# Ephrin-mediated dendrite-dendrite repulsion regulates compartment-specific targeting of dendrites in the central nervous system

**DOI:** 10.1101/2024.10.29.620860

**Authors:** Xiaobing Deng, Sijun Zhu

## Abstract

Neurons often forms synaptic contacts at specific subcellular domains to differentially regulate the activity of target neurons. However, how dendrites are targeted to specific subcellular domains of axons is rarely studied. Here we use *Drosophila* mushroom body out neurons (MBONs) and local dopaminergic neurons (DANs) as a model system to study how dendrites and axons are targeted to specific subcellular domains (compartments) of mushroom body axonal lobes to form synaptic contacts. We found that Ephrin-mediated dendrite-dendrite repulsion between neighboring compartments restricts the projection of MBON dendrites to their specific compartments and prevents the formation of ectopic synaptic connections with DAN axons in neighboring compartments. Meanwhile, DAN neurons in a subset of compartments may also depend on their partner MBONs for projecting their axons to a specific compartment and cover the same territory as their partner MBON dendrites. Our work reveals that compartment-specific targeting of MBON dendrites and DAN axons is regulated in part by a combination of dendrite-dendrite repulsion between neighboring compartments and dendrite-axon interactions within the same compartment.

## INTRODUCTION

One of the fundamental questions in developmental neuroscience is how neurons form precise connections during development. Failure to form proper connections among neurons can lead to functional defects of the nervous system and abnormal behaviors. When presynaptic neurons form synaptic contacts with postsynaptic neurons, synaptic contacts are often formed at specific subcellular domains (Yogev and Shen 2014; Gutman-Wei and Brown 2021). The subcellular specificity of synaptic contacts is functionally important as it could impact neuronal activity differently (Vu and Krasne 1992; Miles et al. 1996).

However, mechanisms that control the subcellular specificity of synaptic contacts are still not fully understood. Most studies investigating mechanisms underlying the subcellular specificity of synaptic contacts have been focused on how presynaptic axons are targeted to distinct subcellular domains of postsynaptic neurons and how cell surface molecules expressed in specific subcellular domains of postsynaptic neurons to attract or repel presynaptic axons to specific subcellular domains of postsynaptic neurons (e.g. (Ango et al. 2004; Suto et al. 2007; Favuzzi et al. 2019; Gutman-Wei and Brown 2021). It has rarely been studied how dendrites of postsynaptic neurons can be guided to specific subcellular domains of presynaptic axons to form synaptic contacts. Further, when different postsynaptic neurons target their dendrites to neighboring subcellular domains of their common presynaptic neurons, whether their subcellular specific targeting is regulated by interactions between dendrites from neighboring subcellular domains is not known.

In the *Drosophila* adult brain, mushroom body (MB) neurons, which are involved in olfactory-associative learning and memory, transmit olfactory and other sensory information to downstream neurons called mushroom body output neurons (MBONs) (Guven-Ozkan and Davis 2014; Cognigni et al. 2018; Aso and Rubin 2020). The later mediate either approaching or avoidance behaviors (Sejourne et al. 2011; Aso et al. 2014b; Owald et al. 2015; Perisse et al. 2016). Meanwhile, local dopaminergic neurons (DANs) that are activated by unconditioned appetitive or aversive stimuli form synaptic contact with both MB neurons and MBONs to modulate synaptic transmission from MB neurons to MBONs and subsequent behavioral outputs, thus inducing memories associated with unconditioned stimuli (Schwaerzel et al. 2003; Schroll et al. 2006; Mao and Davis 2009; Liu et al. 2012; Aso et al. 2014a; Cohn et al. 2015; Yamagata et al. 2015; Takemura et al. 2017). In general, DANs activated by appetitive stimuli attenuate the activity of MBONs mediating avoidance behaviors (Schwaerzel et al. 2003; Schroll et al. 2006; Mao and Davis 2009), whereas DANs activated by aversive stimuli attenuate the activity of MBONs mediating approaching behaviors (Liu et al. 2012; Yamagata et al. 2015). Such specific modulation of MBON activity by DANs relies on precise targeting of MBON dendrites and DAN axons to specific compartments of MB axonal lobes. There are 5 MB axonal lobes, including γ, α’, β’, α, and β lobes, formed by parallel axons from three major types (γ, α’/β’, and α/β) of MB neurons (Fig. 1A) (Lee et al. 1999). These axonal lobes can be divided into 15 compartments, including 5 in the γ lobe, 3 each in the α and α’ lobes, and 2 each in the β and β’ lobes. These compartments abut each other in a non-overlapping manner (Tanaka et al. 2008; Aso et al. 2014a). There are 34 MBONs of 21 types and over 100 DANs of 20 different types. Individual types of MBONs target their dendrites to 1-2 compartments in the MB axonal lobes. Meanwhile, their partner DANs also target their axons to the same compartments such that the activity of individual types of MBONs is only modulated by their partner DANs (Tanaka et al. 2008; Aso et al. 2014a). Such compartment-specific targeting of MBON dendrites and DAN axons provides an excellent model for studying how dendrites and axons are targeted to specific subcellular domains to form synaptic contacts, but very little is known about the mechanisms regulating compartment-specific targeting of MBON dendrites and DAN axons and whether MBON dendrites and DAN axons within the same compartment would interact to regulate each other’s targeting.

**Fig. 1.**
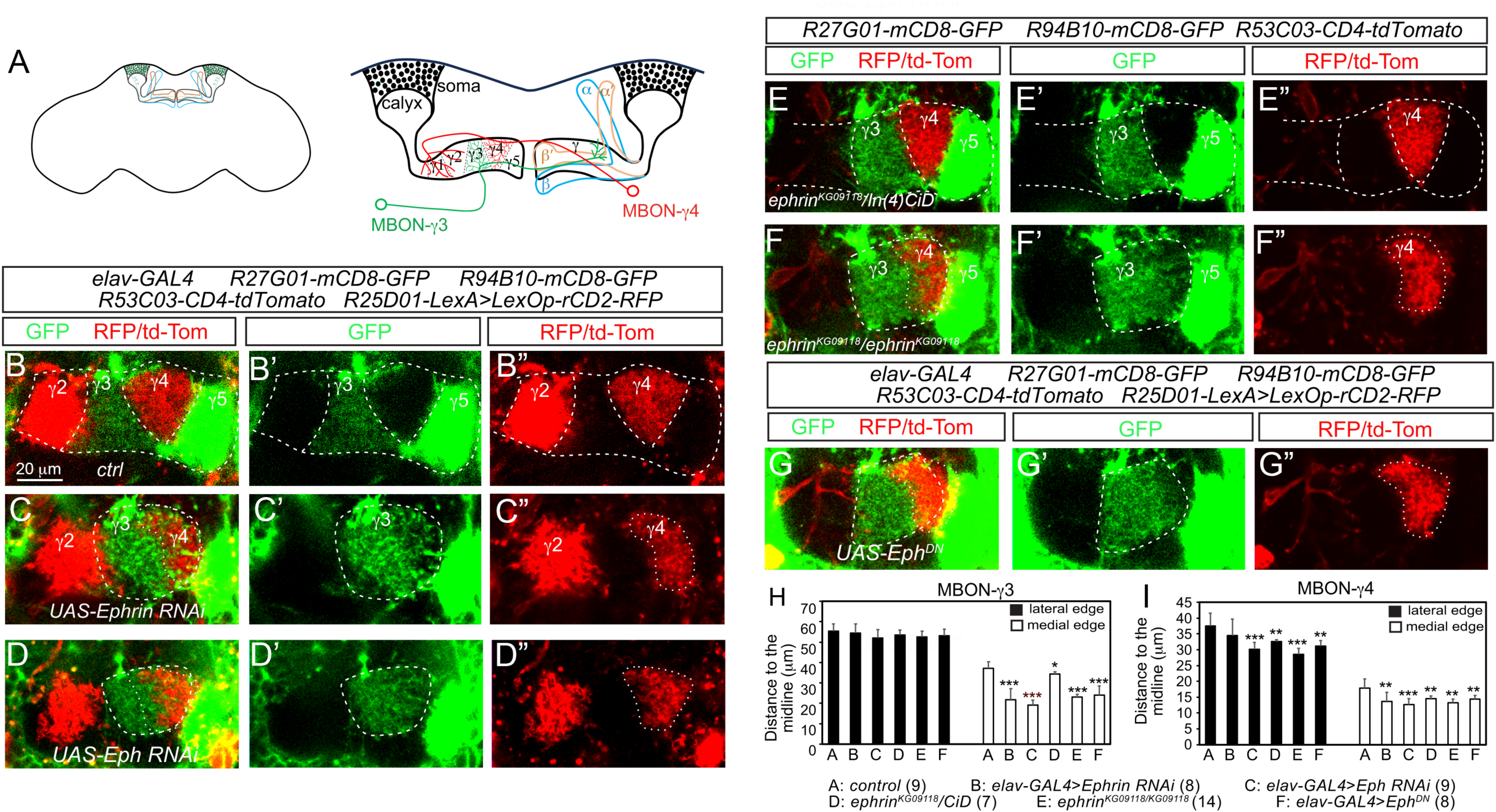
Loss of Ephrin-mediated repulsion leads to defects of compartment-specific target of MBON-γ3 and MBON-γ4 dendrites. In this and all the following figures, images are oriented such that the lateral side of the brain is on the left and the midline of the brain on the right. Only images from one hemisphere are shown. (A) Schematic diagrams of the mushroom body (MB) in the fly brain (left) and an enlarged view of the MB that shows five compartments in the γ lobe and targeting of the dendrites of MBON-γ3 and MBON-γ4 neurons to the γ3 and γ4 compartments (right). (B-B”) Targeting of MBON dendrites labeled with either RFP/tdTom or GFP in γ2 through γ5 compartments in a wild-type adult brain. (C-C”) MBON-γ3 dendrites project into the γ4 compartment and overlap with MBON-γ4 dendrites, which are squeezed into a narrower territory, when Ephrin (B-B”) or Eph (D-D”) is pan-neuronally knocked down. Projection of MBON-γ2 and MBON-γ5 dendrites is not affected by Ephrin or Eph knockdown. (E-E”) Targeting of dendrites of MBON-γ3, MBON-γ4, and MBON-γ5 neurons target their dendrites into their respective compartment in *ephrin^KG09118^/In(4)CiD* heterozygous mutant brains. (F-F”) MBON-γ3 dendrites project into the γ4 compartment, overlapping with MBON-γ4 dendrites, in *ephrin^KG09118^* homozygous mutant brains. (G-G’) Pan-neuronal expression of *UAS-Eph^DN^*results in projection of MBON-γ4 dendrites into the γ4 compartment and overlap between MBON-γ3 dendrites and MBON-γ4 dendrites. (H-I) Quantifications of the distance from the lateral or medial edge of the dendritic territory of MBON-γ3 (H) or MBON-γ4 (I) neurons to the midline of the brain in animals with indicated genotypes. *, *p*<0.05, **, *p*<0.01, \*\*\**p*<0.001, compared to control; student *t*-test. The numbers beside individual genotypes represent sample sizes.

Ephrin is a contact-dependent axon guidance molecule that functions by binding to the Eph receptors (Reber et al. 2007; Klein and Kania 2014). Both Ephrins and Ephs belong to the family of receptor tyrosine kinase (Tuzi and Gullick 1994; Kullander and Klein 2002). Binding of an Ephrin to an Eph receptor could activate bi-directional signaling. It not only activates forward signaling in Eph-expressing cells, but also could activate reverse signaling in Ephrin-expressing cells under certain circumstances (Bruckner et al. 1997; Lu et al. 2001; Mendes et al. 2006; Dudanova et al. 2012). Binding of Ephrins to Ephs mostly promotes cell repulsion (Drescher et al. 1995; Frisen et al. 1998; Rohani et al. 2014), but could also lead to cell adhesion in some cases (Dottori et al. 1998; Davy and Robbins 2000; Huai and Drescher 2001). Ephrins and Eph receptors are well known for their roles in guiding the projection of retinal ganglion axons during retinotectal topographic map formation (Drescher et al. 1995; Frisen et al. 1998; Hornberger et al. 1999; Dearborn et al. 2002; Hindges et al. 2002; Suetterlin and Drescher 2014), but are also involved in regulating axon targting/branching and dendritic patterning of many different types of neurons (Helmbacher et al. 2000; Bossing and Brand 2002; Boyle et al. 2006; Paixao et al. 2013; Anzo et al. 2017). In addition to functioning as axon guidance molecules, Ephrins and Eph receptors are also involved in cell migration, tissue segregation, and cell proliferation, etc. (Xu et al. 1999; North et al. 2009; Senturk et al. 2011; Villar-Cervino et al. 2013).

In our companion paper, we demonstrated that mutual repulsive interactions between neighboring compartments is critical for restricting the targeting of MBON dendrites and DAN axons to their specific compartments and identified Slit that mediates such repulsions in a subset of compartments. Here, we identified another repulsive molecule Ephrin that is expressed in MBON-γ4 neurons to repel of MBON-γ3 dendrites during early pupal stages when MBONs establish their adult-specific dendritic projection patterns and to prevent formation of ectopic synaptic contacts between MBON-γ3 dendrites and PAM-γ4 axons. We further show MBON-γ3 neuron dendrites likely regulate compartment-specific targeting of PAM-γ3 neuron axons either directly or indirectly to ensure that they cover the same territory. Together, results presented in this and our companion paper suggest that compartment-specific targeting of MBON dendrites and DAN axons involves repulsions mediated by different repulsive molecule between neighboring compartments and interactions between MBONs and their partner DANs within the same compartment. These mechanisms act in concert to ensure not only that MBON dendrites and DAN axons not only form synaptic contacts with MBON axons within specific compartments but also that MBON dendrites only form synapses with their partner DAN axons but not these from neighboring compartments.

## RESULTS

### Ephrin-mediated repulsion prevents MBON-γ3 dendrites from expanding into the γ4 compartment

In order to identify additional molecules besides Slit that mediate the repulsive interactions between neighboring compartments in the MB γ lobe, we performed an RNAi knockdown screen of guidance molecules using *elav-GAL4* (Luo et al. 1994) to pan-neuronally express *UAS-RNAi*. Meanwhile, for examining the targeting of MBON dendrites in γ lobe, transgenic lines *R27G01-mCD8-GFP*, *R53C03-CD4-tdTomato*, and *R94B10-mCD8-GFP* we generated (our companion paper) were used to label MBON-γ5β’2a (simplified as MBON-γ5 hereafter), MBON-γ4>γ1,γ2 (simplified as MBON-γ4), and MBON-γ3β’1 (simplified MBON-γ3) neurons, and *LexAop-rCD2-RFP* driven by *R25D01-LexA* to label MBON-γ2γ’1 (simplified as MBON-γ2 hereafter) neurons, respectively. From this screen, we found that knockdown of Ephrin or its receptor Eph led to MBON-γ3 dendrites projecting into the γ4 compartment and extensive overlap with MBON-γ4 dendrites (Fig. 1A-C”). The distance from the medial edge of the dendritic field of MBON-γ3 neurons to the midline of the brain was much reduced, but not the distance from the lateral edge to the midline (Fig. 1G). In contrast, the dendritic field of the MBON-γ4 neurons became narrower with its lateral edge becoming curved instead of being straight as if they were pushed by the invading MBON-γ3 dendrites (Fig. 1A-C”, H). However, targeting of MBON-γ2 and MBON-γ5 dendrites were not affected (Fig. 1A-C”). These phenotypes were further confirmed in *ephrin^KG09118^* mutants or by pan-neuronally expression of a dominant negative form of Eph (Eph^DN^) (Fig. 1D-H). These results suggest that Ephrin regulates compartment-specific targeting of MBON-γ3 and MBON-γ4 dendrites.

### MBON-γ4 neurons express Ephrin to repel MBON-γ3 dendrites

Next we wanted to determine the source of Ephrin. Given that MBON-γ3 dendrites expanded into the γ4 compartment when Ephrin or Eph was knocked and that ablation of MBON-γ4 neurons led to similar albeit weaker phenotypes (our companion paper), it is likely that Ephrin is expressed in MBON-γ4 neurons. Unfortunately, we could not detect Ephrin in MBON-γ4 neurons by immunostaining using the existing Ephrin antibody (Bossing and Brand 2002) or using the GFP trap line *Ephrin^fTRG00598.sfGFP-TVPTBF^* (Sarov et al. 2016) at either pupal or adult stages (data now shown). Therefore, to determine if MBON-γ4 neurons produce Ephrin, we examined if expressing Ephrin in MBON-γ4 neurons would rescue the *ephrin^KG09118^*mutant phenotypes and if knockdown of Ephrin in MBON-γ4 neurons could phenocopy pan-neuronal Ephrin knockdown. Indeed, expressing *UAS-Ephrin-myc* in MBON-γ4 neurons fully rescued the targeting defects of MBON-γ3 and MBON-γ4 dendrites in *ephrin^KG09118^* mutants and fully restored sharp boundary between MBON-γ3 dendrites and MBON-γ4 dendrites(Fig. 2A-C”, E-F). Consistently, knocking down Ephrin in MBON-γ4 neurons resulted in similar phenotypes as pan-neuronal knockdown of Ephrin (Fig. 2D-F). In contrast, knockdown of Ephrin in PAM-γ4, PAM-γ3-5, or MB neurons did not affect the targeting of the dendrites of MBON-γ3 and MBON-γ4 neurons (Supplemental Fig. S1A-E”), nor did expression of *UAS-Ephrin-myc* in PAM-γ4 neurons rescue the phenotypes in *ephrin^KG09118^*mutants (Supplemental Fig. S1C-D”). These results demonstrate that MBON-γ4 neurons express Ephrin to prevent MBON-γ3 neuron dendrites from projecting into the γ4 compartment.

**Fig. 2.**
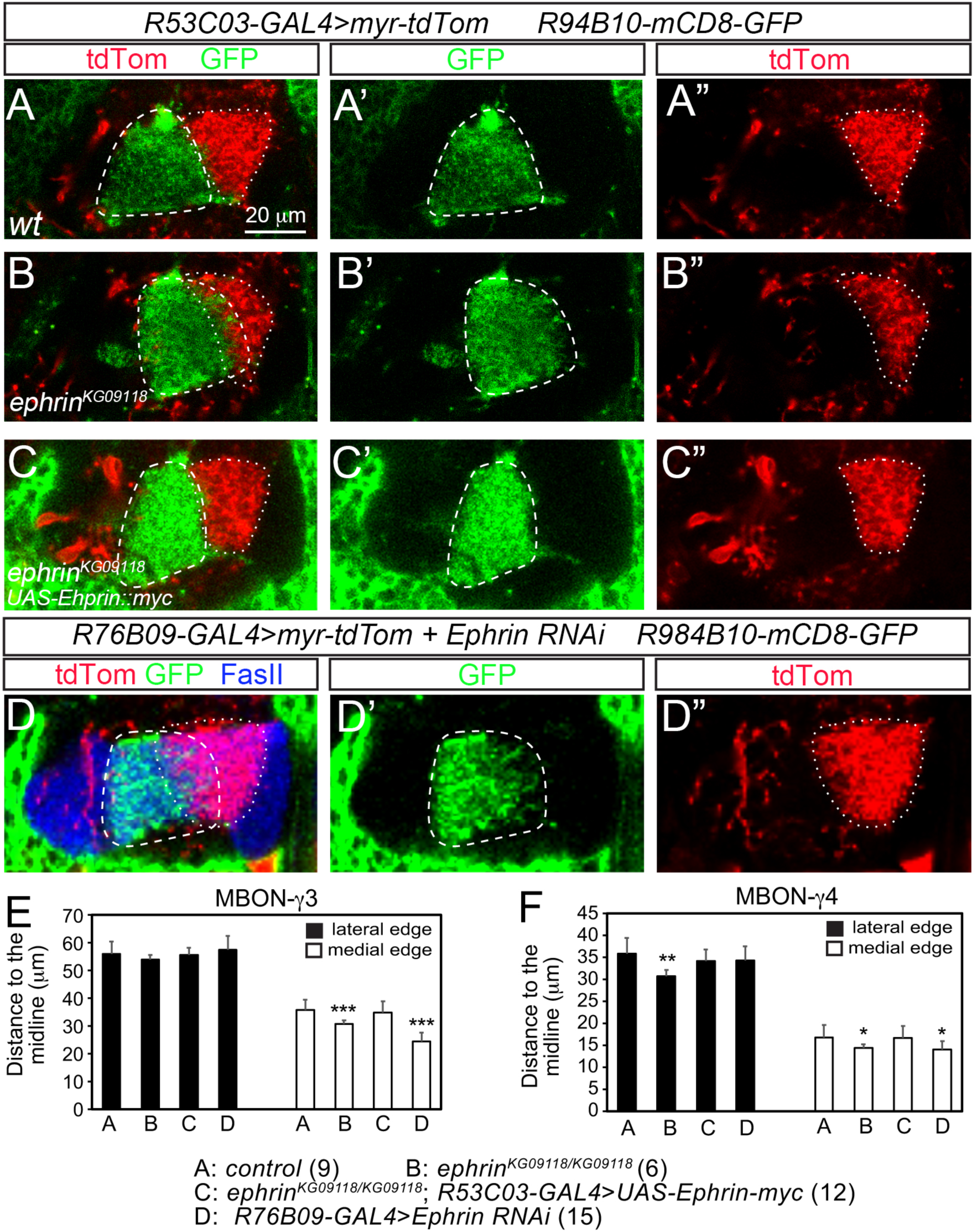
MBON-γ4 neurons likely express Ephrin. In all images, MBON-γ3 dendrites are labeled with GFP and circled with dashed lines and MBON-γ4 dendrites are labeled with tdTom and circled with dotted lines. (A-A’) Projection of MBON-γ3 dendrites and MBON-γ4 dendrites in the γ3 and γ4 compartment in a wild-type brain. (B-B”) MBON-γ3 dendrites project into the γ4 compartment and overlap with MBON-γ4 dendrites in *ephrin^KG09118^* homozygous mutant brains. (C-C”) Expression of *UAS-Ephrin::myc* in MBON-γ4 neurons restores a sharp boundary between MBON-γ3 dendrites and MBON-γ4 dendrites in *ephrin^KG09118^* homozygous mutant brains. (D-D”) MBON-γ3 dendrites project into the γ4 compartment and overlap with MBON-γ4 dendrites when Ephrin is knocked down in MBON-γ4 neurons. (E-F) Quantifications of the distance from the lateral or medial edge of the dendritic territory of MBON-γ3 (E) or MBON-γ4 (F) neurons to the midline of the brain in animals with indicated genotypes. *, *p*<0.05; \*\*\**p*<0.001; compared to control; student *t*-test. The numbers beside individual genotypes represent sample sizes.

### Eph is expressed in MBON-γ3, MBON-γ5, and PAM-γ5 neurons but only knockdown of Eph only affects MBON-γ3 dendrite targeting

Ephrin expressed in MBON-γ4 neurons could directly repel MBON-γ3 dendrites or indirectly by repelling of the PAM-γ3 axons, which may guide the targeting of MBON-γ3 dendrites. However, the lack of targeting defects of MBON-γ3 dendrites after ablation of PAM-γ3 neurons (our companion paper) suggests that it is unlikely that Ephrin repels MBON-γ3 neuron dendrites indirectly. Therefore, MBON-γ3 neuron dendrites probably express Eph and are directly repelled by MBON-γ4 dendrites. Indeed, using the GFP trap line Eph^fTRG00019.sfGFP-TVPTBF^ (simplified as Eph-GFP hereafter) (Sarov et al. 2016) as a reporter, we found that Eph-GFP is expressed in the γ3 compartment and the soma of MBON-γ3 neurons at 2 days APF (Fig. 3A-A”, C-C”). Expression of *UAS-Eph RNAi* in MBON-γ3 neurons driven by two copies of *R94B10-GAL4* largely abolished the expression of Eph-GFP in the γ3 compartment and the soma of MBON-γ3 neurons (Fig. 3B-B”, I), and led to similar phenotypes as pan-neuronal knockdown of Eph (Fig. 3G-H”, J-K). In contrast, expression of *UAS-Eph RNAi* in PAM-γ3 neurons driven by two copies of *R48B04-GAL4* did not affect the expression of Eph-GFP in their soma nor targeting of PAM-γ3 axons (Supplemental Fig. S2A-D”). These results demonstrate Therefore, Ephrin expressed in MBON-γ4 neurons repels MBON-γ3 dendrites directly by binding to Eph expressed in MBON-γ3 neurons. The signal of Eph-GFP in the cell bodies of PAM-γ3 neurons is likely just background.

**Fig. 3.**
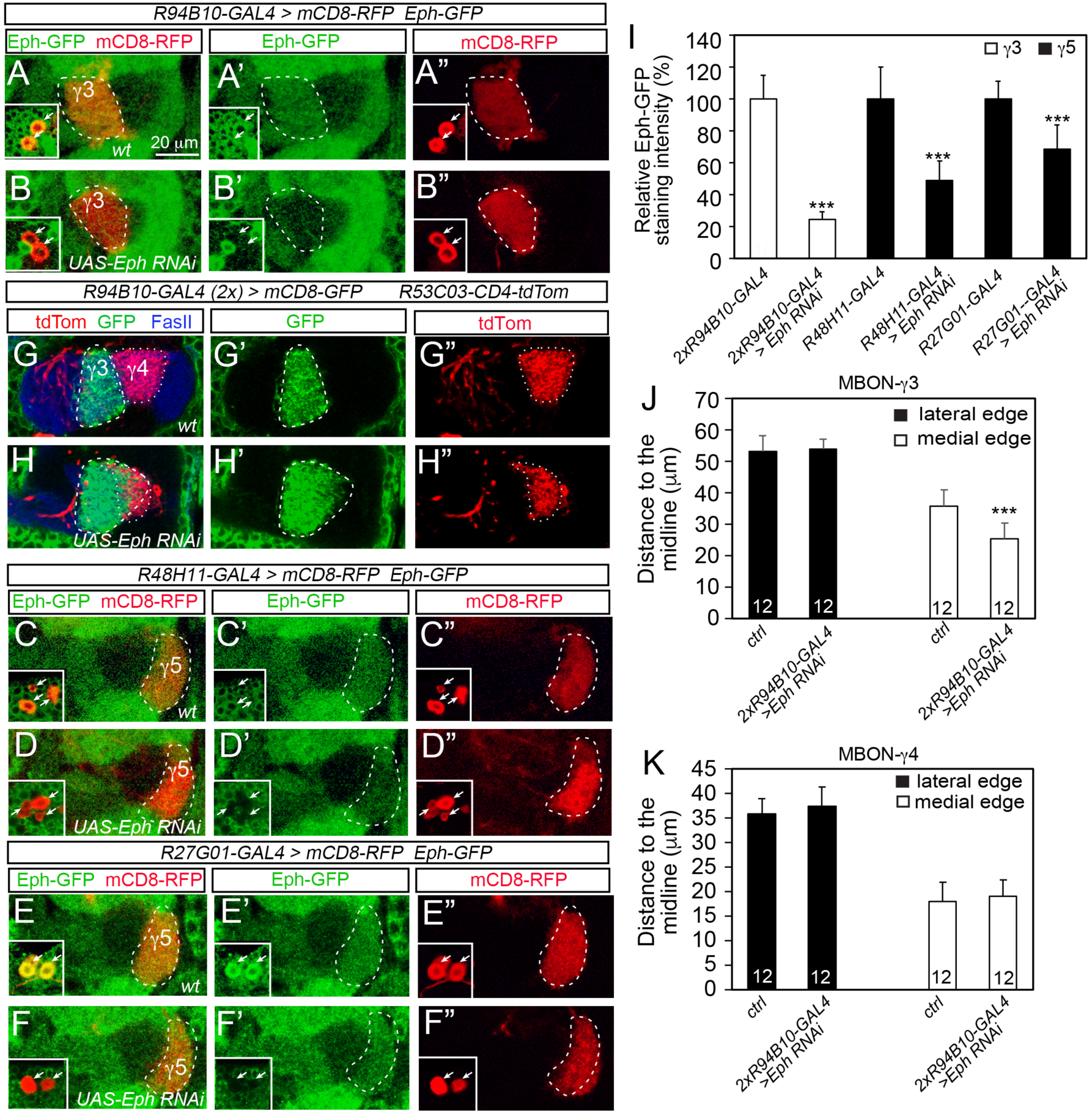
Eph is expressed in MBON-γ3, MBON-γ5, and PAM-γ5 neurons but is only necessary in MBON-γ3 neurons for the compartment-specific targeting of their dendrites. Images in (A-F”) are from brains at 2-3 days APF and images in (G-H”) are from adult brains. (A-A”) Eph-GFP is detected in the γ3 compartment (dashed circles) and in the cell bodies (arrows in insets) of MBON-γ3 neurons, which are labeled with mCD8-RFP. (B-B”) Expression of *UAS-Eph RNAi* in MBON-γ3 neurons leads to near abolishment of Eph-GFP in the cell bodies (arrows in insets) of MBON-γ3 neurons and in the γ3 compartment (dashed circles). (C-C”) Eph-GFP is detected in the γ5 compartment (dashed circles) and in the cell bodies (arrows in insets) of mCD8-RFP-labeled PAM-γ3 neurons. (D-D”) Expression of *UAS-Eph RNAi* in PAM-γ5 neurons results in significant reduction of Eph-GFP expression in the γ5 compartment (dashed circles) and loss of Eph-GFP expression in the cell bodies (arrows in insets) of PAM-γ3 neurons. (E-E”) Eph-GFP is expressed in the γ5 compartment (dashed circles) and in the cell bodies (arrows in insets) of mCD8-RFP-labeled MBON-γ5 neurons. (F-F”) Expression of *UAS-Eph RNAi* in MBON-γ5 neurons results in a similar reduction in Eph-GFP expression in the γ5 compartment (dashed circles) and loss of Eph-GFP in the cell bodies (arrows in insets) of MBON-γ5 neurons labeled with mCD8-RFP. (G-G”) MBON-γ3 dendrites (GFP, dashed circles) and MBON-γ4 dendrites (tdTom, dotted circles) are targeted to the γ3 and γ4 compartment, respectively, without overlapping with each other in a wild-type brain. (H-H”) Knockdown of Eph in MBON-γ3 neurons results in projection of MBON-γ3 dendrites (dashed circles) into the γ4 compartment and overlapping with MBON-γ4 dendrites (dotted circles). (I) Quantifications of relative Eph-GFP staining intensities in the γ3 or γ5 compartment in control brains and brains with Eph knockdown in the MBON-γ3, PAM-γ5, or MBON-γ5 neurons. (J-K) Quantifications of the distance from the lateral or medial edge of the dendritic territory of MBON-γ3 (J) or MBON-γ4 (K) neurons to the midline of the brain in control brains or brains with Eph knockdown in MBON-γ3 neurons. \*\*\**p*<0.001, compared with their corresponding controls; student *t*-test. Numbers in each bar in (I-K) represents sample sizes.

In addition to the γ3 compartment, we found that Eph-GFP was also detected in the γ5 compartment and in the cell bodies of both MBON-γ5 and PAM-γ5 neurons (Fig. 3C-C”, E-E”). Expression of *UAS-Eph RNAi* in MBON-γ5 or PAM-γ5 neurons largely abolished the expression of Eph-GFP in their cell bodies and 30-50% reduction of Eph-GFP in γ5 compartment at 2 days APF (Fig. 3D-D”, F-F”, I) (Fig.3 C-F”, I). However, no targeting defects of MBON-γ5 dendrites and PAM-γ5 neuron axons in the γ5 compartment were observed either at 2 days APF (Fig. 3C-F”) or adult stages (Supplemental Fig. S2E-F”) when Eph was knockdown in these neurons, which is consistent with the results that pan-neuronal knockdown of Ephrin or Eph does not affect targeting of MBON-γ5 dendrites. It is possible that other repulsive molecules expressed in MBON-γ4 and PAM-γ4 neurons function redundantly to repel MBON-γ5 dendrites (and probably also PAM-γ5 axons) given that ablation of MBON-γ4 or PAM-γ4 neurons led to projection of MBON-γ5 dendrites into the γ4 compartment (our companion paper).

### MBON-γ4 neurons extend their dendrites to the γ4 compartment at early pupal stages to repel MBON-γ3 dendrites

To further elucidate how MBON-γ4 dendrites repel MBON-γ3 dendrites, we examined the development of their dendrites at different developmental stages in the wild type or in *ephrin^KG09118^*mutant animals. Since the MB γ lobe undergoes remodeling during early pupal stages (Technau and Heisenberg 1982; Lee et al. 1999), we compared their targeting before, during, and after the remodeling of the γ lobe. At the 3^rd^ instar larval stages, we found MBON-γ3 dendrites and MBON-γ5 dendrites are targeted to neighboring compartments in the medial axonal lobe of larval γ neurons, with the MBON-γ3 dendrites to the shaft of the lobe and MBON-γ5 dendrites to the upper toe and lower toe compartments close the tip of the lobe. There is not gap between MBON-γ3 dendrite and MBON-γ5 dendrites and no MBON-γ4 dendrites were detected between them at the larval stages (Fig. 4A-B). Therefore, MBON-γ4 neurons have not projected their dendrites to the γ lobe yet at this stage. At 18 hours APF, although the larval γ axons have been pruned, the dendrites of MBON-γ3 and MBON-γ5 neurons remain next to each other but are not as dense as at larval stages (Fig. 4C), indicating that their dendrites may also undergo reorganization but are not pruned like the γ lobe. However, MBON-γ4 dendrites are still largely absent from the γ lobe (Fig. 4C). From 20 to 22 hrs APF, MBON-γ4 dendrites start to grow into the γ lobe from the dorsal side between MBON-γ5 dendrites and MBON-γ3 dendrites (Fig. 4D). At 26-28 hrs APF, more MBON-γ4 dendrites have grown into the dorsal half of the γ lobe. Meanwhile, MBON-γ3 dendrites shifted laterally, away from MBON-γ5 dendrites (Fig. 4E). At 36-42 hrs APF when the γ lobe has fully re-extended, MBON-γ4 dendrites have established their adult project patterns and occupied the γ4 compartment, whereas the dendrites of MBON-γ5 and MBON-γ3 neurons have also occupied their respective γ5 and γ3 compartments like in the adult brain (Fig. 4F-G). During the entire process of the projection of MBON-γ4 dendrites into the γ4 compartment, we did not observe overlap of MBON-γ4 dendrites with either MBON-γ3 or MBON-γ5 dendrites (Fig. 4D-G).

**Fig. 4.**
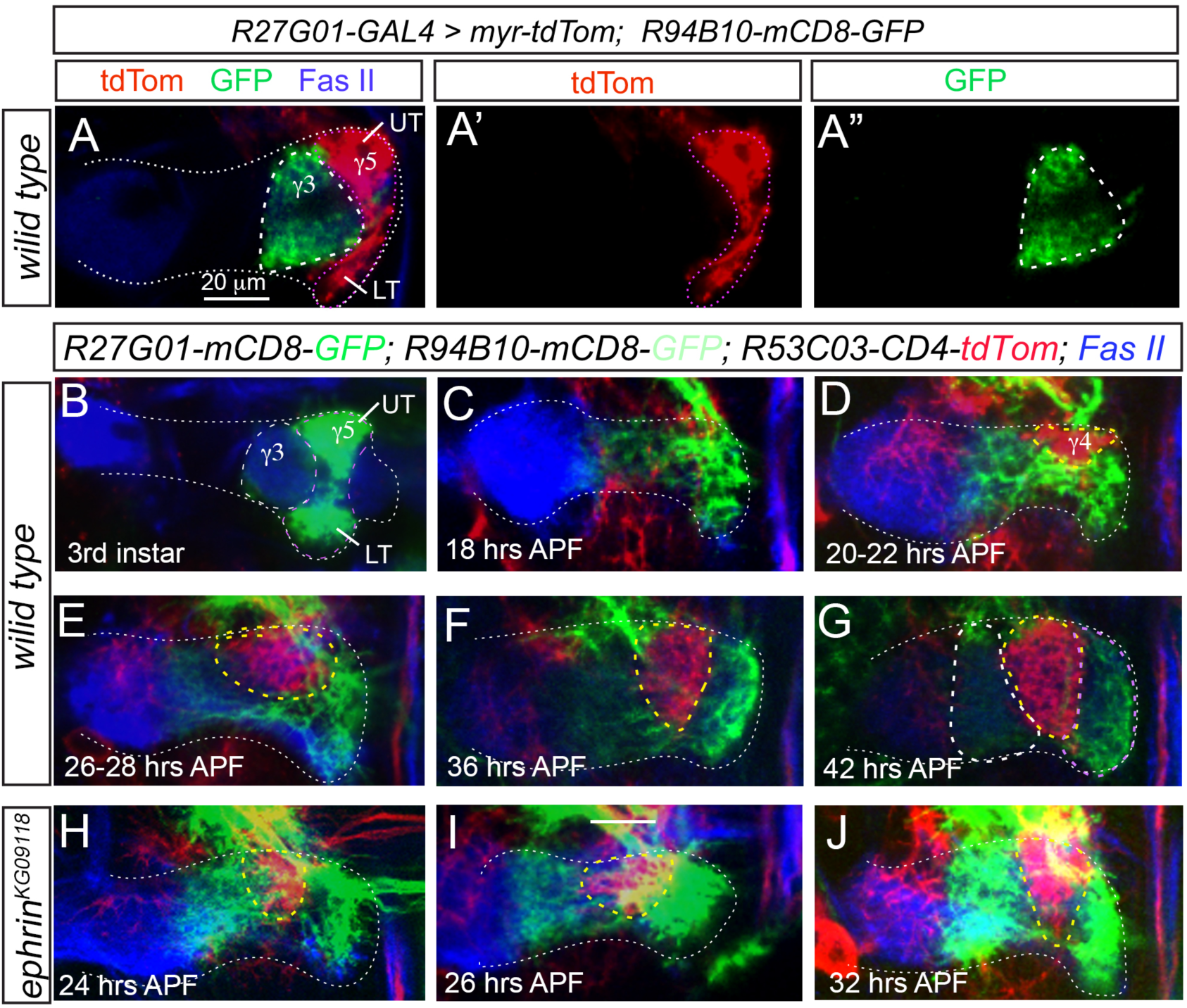
MBON-γ4 neurons project their dendrites to the γ4 compartment during early pupal stages. (A-A”) MBON-γ3 dendrites and MBON-γ5 dendrites occupy neighboring compartments in the medial axonal lobe (Fas II staining in blue, white dotted line) of the larval MB. MBON-γ3 dendrites are labeled with GFP (white dashed circles) and MBON-γ5 dendrites are labeled with tdTom (purple dotted circles). UT, upper toe; LT, lower toe. (B-I) A time-course of the projection of MBON-γ4 dendrites into the γ4 compartment in wild type (B-G) and *ephrin^KG09118^* mutant (H-I) brains during development. MBON-γ4 neuron dendrites are labeled with tdTom (yellow dashed circles), while MBON-γ3 dendrites (white dashed circles) and MBON-γ5 dendrites (purple dashed circles) are labeled with weaker and stronger GFP expression, respectively. In (B), UT, upper toe; LT, lower toe.

In *ephrin^KG09118^*mutants, however, MBON-γ4 dendrites substantially overlapoed with MBON-γ3 dendrites at 24 hrs APF when MBON-γ4 neurons started to project their dendrites into the γ lobe (Fig. 4H). Such overlap continued to exist throughout the rest of development although the majority of MBON-γ3 dendrites gradually shifted laterally (Fig. 4I-J). However, MBON-γ4 dendrites never overlapped with MBON-γ5 neuron dendrites. These results indicate that in *ephrin^KG09118^*mutants, MBON-γ4 dendrites cannot repel MBON-γ3 dendrites when they grow into the region originally occupied by MBON-γ3 dendrites but are repelled from the γ5 compartment.

Together, these results reveal that MBON-γ4 dendrites are targeted to the γ4 compartment only during the regrowth of the γ lobe. The incoming MBON-γ4 dendrites likely push MBON-γ3 dendrites laterally by expressing Ephrin. Meanwhile they are repelled from the γ5 compartment by Slit expressed in PAM-γ5 neurons (see our companion paper).

### Remodeling is not required for the compartment-specific targeting of MBON dendrites in the γ lobe

Since MBON-γ4 dendrites are only targeted to the γ lobe during the regrowth of the γ lobe and it seems that MBON-γ3 dendrites undergo reorganization during early pupal stages, we asked whether remodeling of the γ lobe is essential for establishing the adult projection patterns of MBONs in the γ lobe and if MBON-γ3 and MBON-γ4 neurons involves ecdysone signaling-mediated remodeling for establishing their adult elaboration pattern. To answer this question, we expressed the dominant negative form of eIF4A (elF4A^DN^) in MB γ neurons and a dominant-negative form of ecdysone receptor B1 (EcRB1^DN^) or *UAS-EcRB1 RNAi* in MBON-γ3 or MBON-γ4 neurons. eIF4A is required for pruning of MB γ neuron axon and EcRB1 is ubiquitously required for neuronal remodeling during early pupal stages (Lee et al. 2000; Rode et al. 2018). However, we did not observe obviously changes in the compartment-specific targeting of the dendrites of MBON-γ3, MBON-γ4, and MBON-γ5 neurons after blocking the pruning of the γ lobe (Supplemental Fig. S3A) or ecdysone signaling in MBON-γ3 or MBON-γ4 neurons (Supplemental Fig. S3B-C), indicating that neither remodeling of the γ lobe nor ecdysone signaling is required for establishing adult-specific compartment-specific targeting of their dendrites.

### PAM-γ3 axons may in part depend on MBON-γ3 dendrites for their compartment-specific targeting

In each compartment of MB axonal lobes, MBON dendrites and their partner DAN axons share the same territory (Aso et al. 2014a). How do MBON dendrites and their partner DAN axons ensure to cover the same territory? Are there any direct or indirect interactions between them or are they guided to the same compartment independently? To address this question, we next examined how defects in targeting of MBON dendrites or ablation of MBONs would affect the targeting of DAN axons or vice versa. We focused on MBONs and DANs in the γ3 and γ4 compartment for detailed analyses.

In the γ3 compartment, we found that when the dendritic field of MBON-γ3 neurons expanded medially due to knockdown of Eph, PAM-γ3 axons also expanded medially and remained completely overlapping with MBON-γ3 dendrites (Fig. 5A-B’, H). Whereas when MBON-γ3 neurons were ablated, PAM-γ3 axons shrank into a narrow strip, sandwiched between the expanded MBON-γ4 dendrites (also see our companion paper) and the γ2 compartment, although they were still targeted to the γ lobe (Fig. 5D-D’, H). In contrast, when Robo3 was knocked down in PAM-γ3 neurons, which led to projection of their axons into the γ2 compartment (our companion paper), MBON-γ3 dendrites remained in the γ3 compartment and did not follow the expanded PAM-γ3 axons (Fig. 5C-C’, I). This is consistent with the results that ablation of PAM-γ3 neurons did not affect the targeting of MBON-γ3 dendrites (our companion paper). Therefore, PAM-γ3 neurons may partly depend on MBON-γ3 neurons for projecting their axons to the γ3 compartment and cover the same territory as MBON-γ3 dendrites but not the other way around. Meanwhile, the γ lobe may express an attractive molecule to guide the projection of PAM-γ3 axons to the γ lobe as PAM-γ3 axons are still targeted to the γ lobe when MBON-γ3 neurons were ablated. In support of the the potential role of MBON-γ3 dendrites in guiding PAM-γ3 axons, we found that PAM-γ3 neurons project their axons to the γ3 compartment after MBON-γ3 dendrites. Only after 22 and 24 hrs APF did PAM-γ3 neurons start to project their axons into the area where MBON-γ3 neuron dendrites are but not at the 3rd instar and 0hrs APF (Supplemental Fig. S4), which is consistent with recent reports that PAM-γ3 neurons are born post-embryonically and they are adult-specific MB input neurons (Lee et al. 2020; Truman et al. 2023). MBON-γ3 neurons may regulate the compartment-specific targeting of DAN-γ3 axons directly by expressing an attractive molecule to guide the projection of DAN-γ3 axons, indirectly by repelling MBON-γ4 dendrites, which may in turn repel DAN-γ3 axons, or both (see discussion below).

**Fig. 5.**
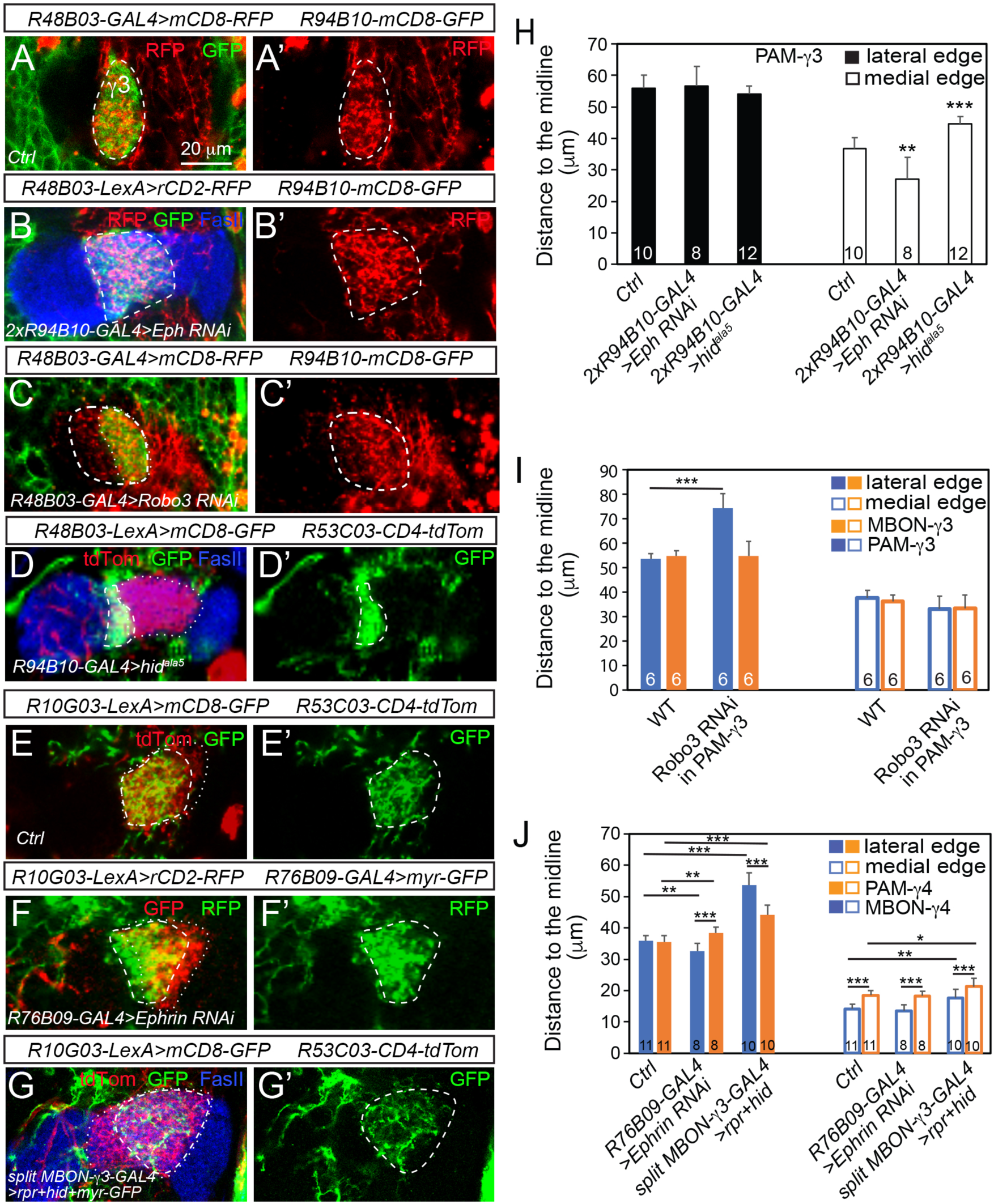
Potential attractive interactions exist between MBON-γ3 dendrites and PAM-γ3 axons but not between MBON-γ4 dendrites and PAM-γ4 axons. (A-A’) PAM-γ3 axons (RFP) completely overlap with MBON-γ3 dendrites (GFP) in the γ3 compartment (dashed circles) in a wild-type adult brain. (B-B’) The of PAM-γ3 axonal field also expands medially and covers the same territory (dashed circles) as the medially expanded MBON-γ3 dendritic field when Eph is knocked down in MBON-γ3 neurons. The MB γ lobe is labeled with Fas II staining in blue. (C-C’) Knockdown of Robo3 in PAM-γ3 neurons only leads lateral expansion of the axonal field (dashed circles) of PAM-γ3 neurons but not the dendritic field of MBON-γ3 neurons. (D-D’) Genetic ablation of MBON-γ3 neurons leads to shrinkage of the axonal field (dashed circles) of PAM-γ3 neurons and expansion of the dendritic field (dotted circles) of MBON-γ4 neurons. The MB γ lobe is labeled with Fas II staining in blue. (E-E’) In a wild-type adult brain, the axonal field (dashed circles) of PAM-γ4 neurons (GFP in green) shares the same lateral edge with the dendritic field (dotted circles) of MBON-γ4 neurons (tdTom in red), but the medial edge of PAM-γ4 axonal field is slightly further from the midline than that of the dendritic field of MBON-γ4 neurons. (F-F’) Knockdown of Ephrin in MBON-γ4 neurons (GFP in red) leads to shifting of the lateral edge of the MBON-γ4 dendritic field (dotted circles) medially but does not affect the targeting of axons of PAM-γ4 neurons (RFP in green, dashed circles). (G-G’) Genetic ablation of MBON-γ3 neurons results in dramatic lateral expansion of the dendritic field of MBON-γ4 neurons (tdTom in red, dotted circles) but only minimal lateral shift of the axonal field of PAM-γ4 neurons (GFP in green, dashed circles). (H-J) Quantifications of the distance from the midline to the lateral or medial edge of the axonal field of PAM-γ3 (H, I) or PAM-g4 neurons (J), or the dendritic field of MBON-γ3 (I) or MBON-γ4 (J) neurons in animal with indicated genotypes. *, *p*<0.05, **, *p*<0.01, \*\*\**p*<0.001; student *t*-test. The numbers on individual bars represent sample sizes.

In contrast to PAM-γ3 and MBON-γ3 neurons, we found that MBON-γ4 dendrites and PAM-γ4 axons are targeted to the γ4 compartment independently. In the wild type brains, PAM-γ4 axons and MBON-γ4 dendrites share the same territory, and their lateral edges exactly overlap (Fig. 5E-E’, J). However, when the lateral edge of the dendritic field of the MBON-γ4 neurons shifted medially as a result of Ephrin or Eph knockdown, the lateral edge of the axonal field of PAM-γ4 neurons barely shifted and no longer overlapped with the lateral edge of MBON-γ4 dendrites (Fig. 5F-F’, J). As a result, PAM-γ4 axons extensively overlapped with expanded MBON-γ3 dendrites rather than being pushed into a narrower territory like MBON-γ4 dendrites (Fig. 6A-B”). Similarly, when the dendritic field of MBON-γ4 neurons expanded laterally due to ablation of MBON-γ3 neurons, PAM-γ4 axons did not follow (Fig. 5G-G’, J). On the other hand, ablation of PAM-γ4 neurons did not affect the targeting of MBON-γ4 dendrites (our companion paper). Therefore, although PAM-γ4 axons and MBON-γ4 dendrites share the same territory, they are likely targeted to the γ4 compartment independently. The lack of expansion or contraction of the axonal field of PAM-γ4 neurons after ablation of MBON-γ3 neurons or knockdown of Ephrin in MBON-γ3 neurons also indicates that unlike MBON-γ4 neuron dendrites, PAM-γ4 axons are not repelled by MBON-γ3 neuron dendrites either directly or indirectly.

**Fig. 6.**
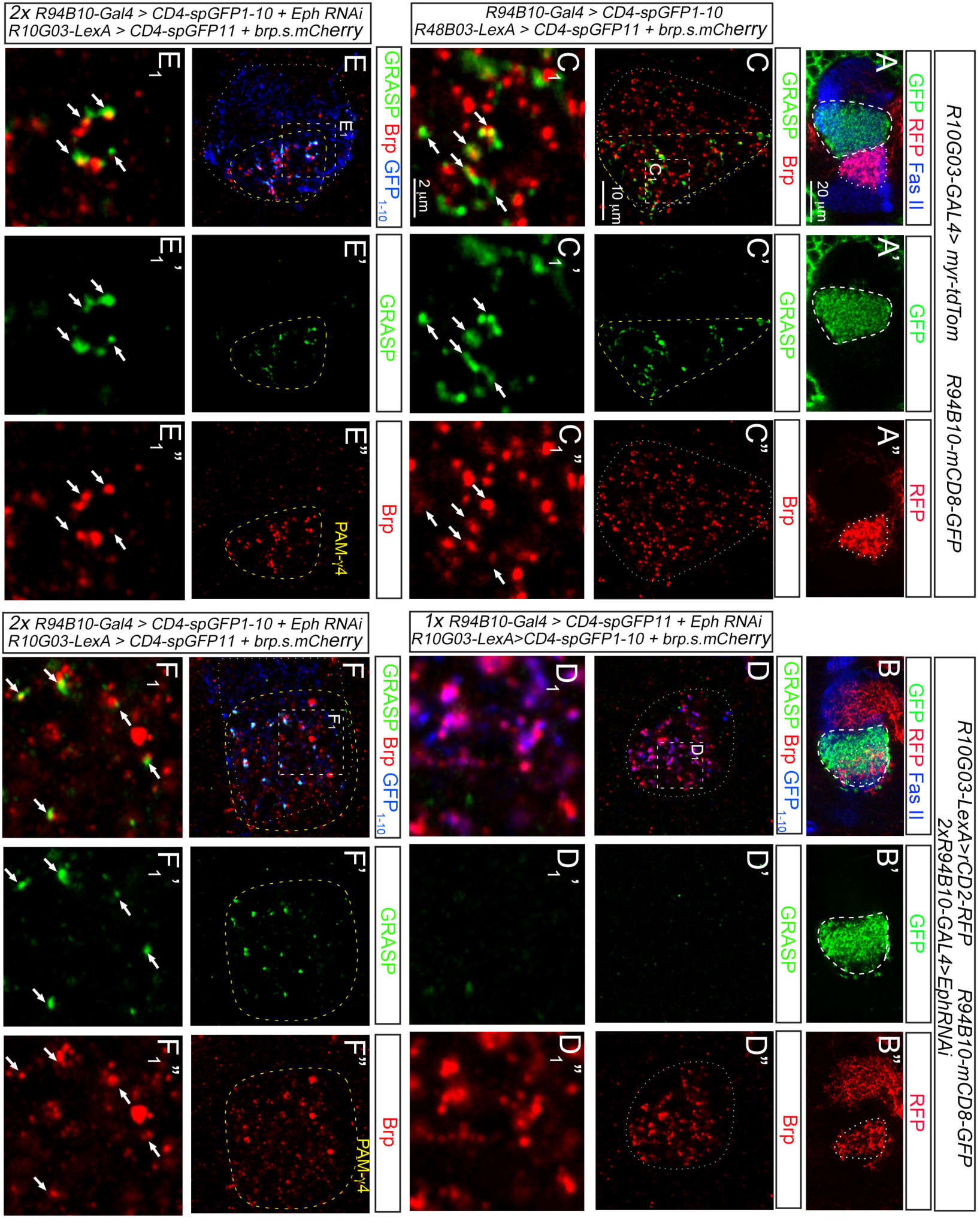
Knockdown of Eph in MBON-γ3 neurons results in ectopic synaptic formation between MBON-γ3 dendrites and PAM-γ4 axons. In (A-B”), MBON-γ3 dendrites are labeled with GFP and outlined with dashed circles and PAM-g4 axons are labeled with RFP and outlined with dotted circles. (A-A”) MBON-γ3 dendrites and PAM-γ4 axons occupy two neighboring compartments without overlapping with each other in wild-type brains. (B-B”) MBON-γ3 dendrites overlap with PAM-γ4 axons when Eph is knocked down in MBON-γ3 neurons. (C-C_1_”) Synaptic contacts are detected between MBON-γ3 dendrites and PAM-γ3 axons with GRASP in wild-type brains. Dotted circles outline PAM-γ3 axonal field. Dashed lines outline the region where GRASP signals are detected. (C_1_-C_1_”) are an enlarged view of the C_1_ area (dashed square) indicated in (C). Arrows point to GRASP signals that overlap with the presynaptic marker Brps.mCherry. (D-D”) No GRASP signals are detected between MBON-γ3 dendrites and PAM-γ4 axons when *UAS-Eph RNAi* is driven by one copy of *R94B10-GAL4*. Dotted circles outline the axonal field of PAM-γ4 neurons. (D_1_-D_1_”) are an enlarged view of the D_1_ area (dashed squares) indicated in (D). (E-F1”) GRASP signals are detected between MBON-γ3 dendrites and PAM-γ4 axons when *UAS-Eph RNAi* is driven by two copies of *R94B10-GAL4*. (E-E_1_”) and (F-F_1_”) are images from two different brains. Dotted circles outline the dendritic field of MBON-γ3 neurons and dashed circles outline the axonal field of PAM-γ4 neurons. (E_1_-E_1_”) and (F_1_-F_1_”) are enlarged views of the E_1_ and F_1_ areas (dashed squares) indicated in (E) and (F), respectively. Arrows point to GRASP signals that overlap with the presynaptic marker Brp.s.mCherry.

### Defects in targeting of MBON dendrites result in ectopic synaptic contacts between MBONs and DANs from neighboring compartments

MBON dendrites normally form synaptic contacts with their partner DAN axons within the same compartment, but not with DAN axons targeted to neighboring compartments. The defects of compartment-specific targeting MBON-γ3 dendrites and extensive overlap between MBON-γ3 dendrites and PAM-γ4 axons resulting from the loss of Ephrin-mediated repulsion led us to ask whether any ectopic synaptic contacts could be formed between MBON-γ3 neuron dendrites and PAM-γ4 axons. To answer this question, we employed the GFP Reconstitution Across *Synaptic* Partners (*GRASP*) technology (Feinberg et al. 2008), which has been used extensively to detect synaptic contacts between neurons (Gorostiza et al. 2014; Yoshino et al. 2017; Zhao et al. 2022). Using this approach, we could successfully detect synaptic contacts between MBON-γ3 dendrites and PAM-γ3 axons in wild type brains as indicated by reconstituted GFP signals that overlapped with the presynaptic marker Bruchpilot (Brp) tagged with mCherry (Brp-mCherry) (Fig. 6C-C_1_”), but not between MBON-γ3 dendrites and PAM-γ4 axons when only one copy of when only one copy of *R94B10-GAL4* was used to drive the expression of *UAS-Eph RNAi*, which did not cause targeting defects of the dendrites of MBON-γ3 and MBON-γ4 neurons (Fig. 6D-D_1_”). However, when Eph was knocked down with two copies of *R94B10-GAL4* in MBON-γ3 neurons, which led to extensive overlap between MBON-γ3 dendrites and PAM-γ4 axons, we detected synaptic contacts between them as indicated by positive reconstituted GFP signals colocalized Brp-mCherry (Fig. 6E-F_1_”). Therefore, defects in compartment-specific targetring of MBON dendrites could lead to formation of ectopic synaptic contacts, which could potentially lead to aberrant synaptic transmission between MBON neurons and DAN neurons and subsequence behavioral outputs.

## DISCUSSION

In this study, we identify Ephrin as a repulsive molecule that regulates compartment-specific targeting of MBON-γ3 dendrites by mediating the repulsion between MBON-γ4 dendrites and MBON-γ3 dendrites. Together with our finding that Slit mediates the repulsion of MBON dendrites and DAN axons in the γ3 and γ4 compartments by PPL1-γ2 and PAM-γ5 axons reported in our companion paper, our work demonstrates that to restrict the projection of MBON dendrites within their specific compartments requires repulsion from neighboring compartments on both sides, which can be mediated by different molecules and different types of cells (Fig.7). Further, in some compartments like γ3, compartment-specific targeting of DAN axons (or MBON dendrites) might be also regulated either directly or indirectly by their partner MBON-γ3 dendrites (or DAN axons) to ensure they cover the same territory.

**Fig. 7.**
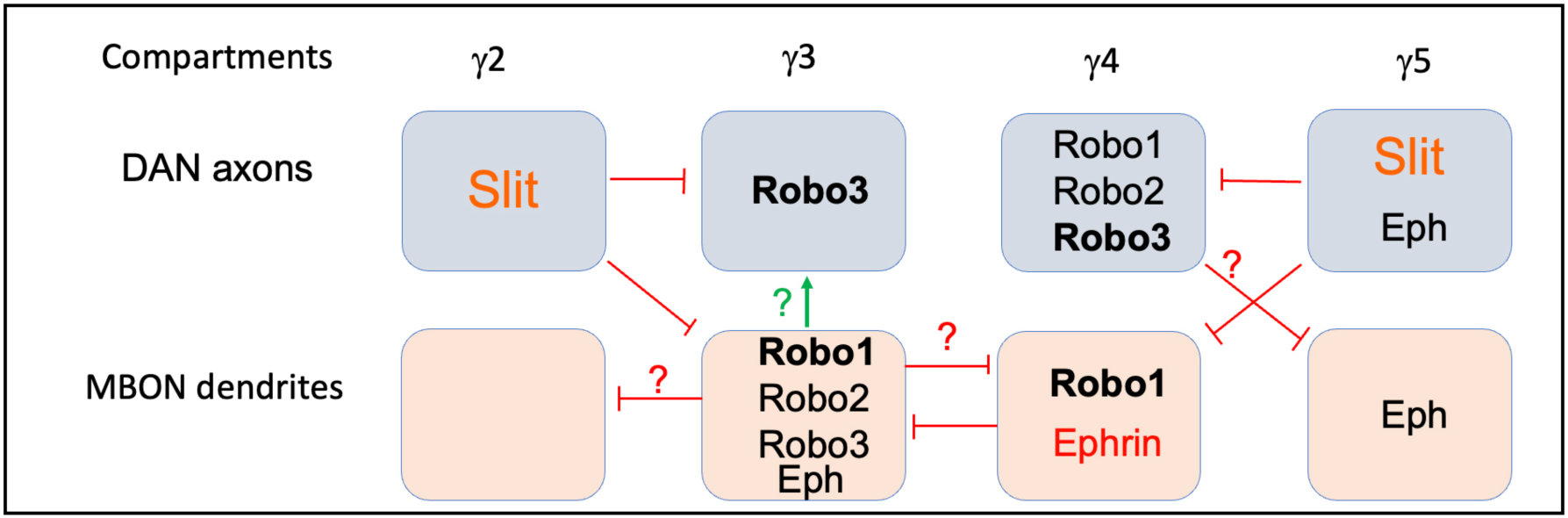
A working model of the regulation of compartment-specific targeting of MBON dendrites and DAN axons by Slit and Ephrin-mediated repulsion. Slit is expressed in PPL1-γ2 axons and PAM-γ5 axons to repel dendrites of MBON-γ3 and MBON-γ4 neurons and axons of PAM-γ3 and PAM-γ4 neurons by binding to different Robo receptors. Ephrin is expressed in MBON-γ4 neurons to prevent MBON-γ3 dendrites from projecting into the γ4 compartment. Eph is also expressed in MBON-γ5 and PAM-γ5 neurons but does not mediate the repulsion of their dendrites or axons by Ephrin.

Ephrin is one of the classic axon guidance molecules. It functions mainly as a contact-dependent repellent and has been well studied in the formation of retinotectal topographic map (Drescher et al. 1995; Frisen et al. 1998; Dearborn et al. 2002; Hindges et al. 2002; Suetterlin and Drescher 2014). In *Drosophila*, Ephrin has also been shown to restrict the growth of axonal branches of photoreceptors within specific columns in the optic lobe and dendrites of projection neurons to specific glomeruli in the antennal lobe (Anzo et al. 2017; Liu et al. 2017). Although we are not able to detect the expression of endogenous Ephrin, our data from Ephrin knockdown and rescue of *ephrin* mutant phenotypes clearly indicate that MBON-γ4 neurons are the source of Ephrin, which is also consistent with the phenotype resulting from ablation of MBON-γ4 neurons (our companion paper). MBON-γ4 neurons express Ephrin to push away dendrites of MBON-γ3 neurons while they project their dendrites into the γ4 compartment during early pupal stages. Projection of MBON-γ4 dendrites is likely guided by an attractive molecule expressed in the MB γ lobe. Since ablation of PAM-γ4 and PAM-γ3 neurons also leads to expansion of MBON-γ3 neuron dendrites into the γ4 compartment (our companion paper), a repulsive molecule expressed in PAM-γ4 neurons may act together with Ephrin to prevent the projection of MBON-γ3 neuron dendrites into the γ4 compartment. On the other hand, given that the dendritic field of MBON-γ4 neurons became narrower when MBON-γ3 dendrites invaded into the γ4 compartment but expanded laterally when MBON-γ3 neurons were ablated (our companion paper), MBON-γ3 neurons may also express a repulsive cue to repel MBON-γ4 dendrites. A recent study reported that Sema1a is expressed in MBON-γ3 neurons (Lin et al. 2022). However, knockdown of Sema1a in MBON-γ3 neurons does not affect the targeting of MBON-γ4 dendrites (our unpublished data). Therefore, MBON-γ3 neurons may express other repulsive molecules to repel MBON-γ4 dendrites.

Surprisingly, although Eph is also expressed in MBON-γ5 and PAM-γ5 neurons, the compartment-specific targeting of their dendrites and axons does not seem to be affected by either Ephrin or Eph knockdown. However, since ablation of MBON-γ4 neurons leads to projection of MBON-γ5 dendrites into the γ4 compartment, MBON-γ4 neurons may express another repulsive molecule that function redundantly with Ephrin to repel MBON-γ5 dendrites and PAM-γ5 axons. Further, given that ablation of PAM-γ4 neurons also results in projection of MBON-γ5 dendrites into the γ4 compartment, both MBON-γ4 and PAM-γ4 neurons are required to prevent MBON-γ5 dendrites (and probably also the PAM-γ5 axons) from projecting into the γ4 compartment, but PAM-γ4 neurons may play a more prominent role because ablation of PAM-γ4 neurons leads to more MBON-γ5 dendrites projecting into the γ4 compartment. To fully understand how MBON-γ5 dendrites and PAM-γ5 axons are repelled from the γ4 compartment, it would be essential to identify other repulsive molecules expressed in MBON-γ4 and PAM-γ4 neurons in the future.

In addition to mutual repulsive interactions between neighboring compartments, our work demonstrates that compartment-specific targeting of the axons of PAM-γ3 neurons also depends on their partner MBON-γ3 neurons. MBON-γ3 neurons could regulate the targeting of PAM-γ3 axons directly by providing an attractive cue to guide the projection of PAM-γ3 axons. Alternatively, MBON-γ3 neurons could regulate the targeting of PAM-γ3 axons indirectly by repelling MBON-γ4 dendrites, which may also repel PAM-γ3 axons. The direct mechanism could explain why PAM-γ3 axons and MBON-γ3 dendrites always cover the same territory no matter whether in the wild type brains or when MBON-γ3 dendrites expanded due to knockdown of Ephrin, but it could not explain why the axonal field of PAM-γ3 neurons would become a narrow strip when MBON-γ3 neurons were ablated. In contrast, the indirect mechanism could explain all these scenarios. When MBON-γ3 neurons were ablated, MBON-γ4 dendrites expanded laterally and pushed PAM-γ3 axons laterally through repulsion. Meanwhile, PAM-γ3 axons are also repelled by Slit secreted from PPL1-γ2 neurons (our companion paper). Therefore, PAM-γ3 axons would be sandwitched between the expanded MBON-γ4 dendrites and PPL1-γ2 axons, occupying a narrow strip.

Whereas in wild type animals or when MBON-γ3 dendrites expanded into the γ4 compartment due to knockdown of Ephrin, MBON-γ3 dendrites push away MBON-γ4 dendrites, thus allowing PAM-γ3 axons to grow into the area occupied by MBON-γ3 dendrites. Although PAM-γ3 axons do not seem to be repelled by Ephrin expressed in MBON-γ4 neurons, as discussed above MBON-γ4 neurons may express another repulsive molecule to repel PAM-γ3 axons. Therefore, we favor the indirect mechanism, but do not rule out the direct mechanism. It is also possible that these two mechanisms work together to ensure that PAM-γ3 axons and MBON-γ3 dendrites cover the same territory.

However, the dependency of DAN axons on MBON dendrites for compartment-specific target does not happen in all compartments. At least in the γ4 compartment, we found that shifting or expansion of the dendritic field of MBON-γ4 neurons had minimal effects on the projection of PAM-γ4 axons. A recent study also shows that in the absence of Dpr12, which leads to failure of projection of PAM-γ4 axons to the γ lobe and failure of extension of the MB γ lobe beyond the γ3 compartment, MBON-γ4 dendrites are still targeted to the shortened MB γ lobes (Bornstein et al. 2021). Therefore, MBON-γ4 dendrites and PAM-γ4 axons are guided to the γ lobe independently by distinct attractive molecules. Since blocking the remodeling of the γ lobe does not affect the targeting of MBON-γ4 dendrites, the expression of the attractive molecule that guide MBON-γ4 dendrites may not depend on the remodeling of the γ lobe. In line with this notion, Dpr12 that guides the projection of PAM-γ4 axons during early pupal stage is also expressed in the medial lobe of larval MBs (Bornstein et al. 2021). However, these attractive molecules may only guide the projection of MBON-γ4 dendrites and PAM-γ4 axons to the γ lobe but not to a specific compartment. To ensure they are targeted to the same compartment and cover the same territory, they have to rely on other shared mechanisms such as repulsions by both PAM-γ5 axons and MBON-γ3 dendrites.

Finally, our work suggests that compartment-specific targeting of MBON dendrites and DAN axons is critical for ensuring that MBON dendrites only form synaptic contacts with their partner DAN axons but not those from neighboring compartments in addition to forming synapses with MB axons. Our GRASP results show that when MBON-γ3 dendrites project to the γ4 compartment due to the loss of Ephrin-mediated repulsion, they could form synaptic contacts with PAM-γ4 axons. These results, together with the similar results that knockdown of Robo receptors in MBON-γ3 neurons leads to formation of ectopic synaptic connections between MBON-γ3 dendrites and PPL1-γ2 axons (our companion paper), suggest that MBON neurons have ability to form synaptic contacts with DAN axons from neighboring compartments, but compartment-specific targeting of MBON dendrites and DAN axons ensures that MBON dendrites only form synapses with their partner DAN axons.

## MATERIALS AND METHODS

### Fly husbandry and strains

For examining the function of Ephrin in compartment-specific targeting of MBONs and DANs, we used *UAS-ephrin RNAi* (P{TRiP.HMS01289}attP2) (Bloomington *Drosophila* Stock Center [BDSC] #34614), *ephrin^KG09118^*(Boyle et al. 2006) (BDSC #15162) for Ephrin RNAi knockdown and mutant phenotypic analyses, respectively. *UAS-ephrin::myc* (Boyle et al. 2006) (a gift from Thomas JB) was used to rescue ephrin mutant phenotypes. For functional analyses of Eph, *UAS-eph RNAi* lines (P{TRiP.HMS04998}attP40, BDSC #60006, and P{TRiP.GL00192}attP2)attP2, BDSC #35290) were used for RNAi knockdown of Eph and *UAS-Eph^DN^*line (Dearborn et al. 2002) (a gift from Kunes S.) was used for dominant negative inhibition of Eph. A BAC transgenic line expressing Eph::GFP (PBac{fTRG00019.sfGFP-TVPTBF}VK00033, Vienna *Drosophila* Resource Center [VDRC] #v318008) was used for examining the expression of endogenous Eph. For inhibiting neuronal remodeling, *UAS-eIF4A^E172Q^*(Rode et al. 2018) (a gift from S. Rumpf), *UAS-EcRB1^DN^*(Cherbas et al. 2003) (BDSC #6869) or *UAS-EcRB1 RNAi* (BDSC #58286) were expressed in the MB γ neurons, MBON-γ3 neurons, or MBON-γ4 neurons. *UAS-hid^ala5^*(Bergmann et al. 2002) or *UAS-hid* together with *UAS-reaper (rpr)* (Zhou et al. 1997) were used for genetic ablation of neurons. For the genetic ablation and RNAi knockdown, animals were raised at 29°C to enhance the expression of UAS-transgenes. For GRASP analyses, *UAS-CD4-spGFP_1-10_*, *LexAop-CD4-SpGFP_11_* (Gordon and Scott 2009), and *LexAop-nSyp-spGFP_1-10_* (Macpherson et al. 2015) were expressed in post- and pre-synaptic neurons. *LexAop-brp.s.mCherry* (Berger-Muller et al. 2013) was co-expressed with *LexAop-CD4-SpGFP_11_* to mark presynaptic sites. *R27G01-mCD8-GFP*, *R94B10-mCD8-GFP*, *R53C03-CD4-tdTomato* (see our companion paper) were used for labeling of MBON-γ5, MBON-γ3 and MBON-γ4 neurons. Expression of *LexAop-rCD2-RFP*, *UAS-myr-tdTomato*, *UAS-mCD8-RFP*, *UAS-myr-GFP*, *UAS-rCD2-RFP*, and *UAS-mCD8-GFP* driven by cell-type specific LexA or GAL4 lines were used to label neurons of interest. GAL4 and LexA lines used for driving UAS/LexAop-transgene expression are listed below in Table 1.

**Table 1.**
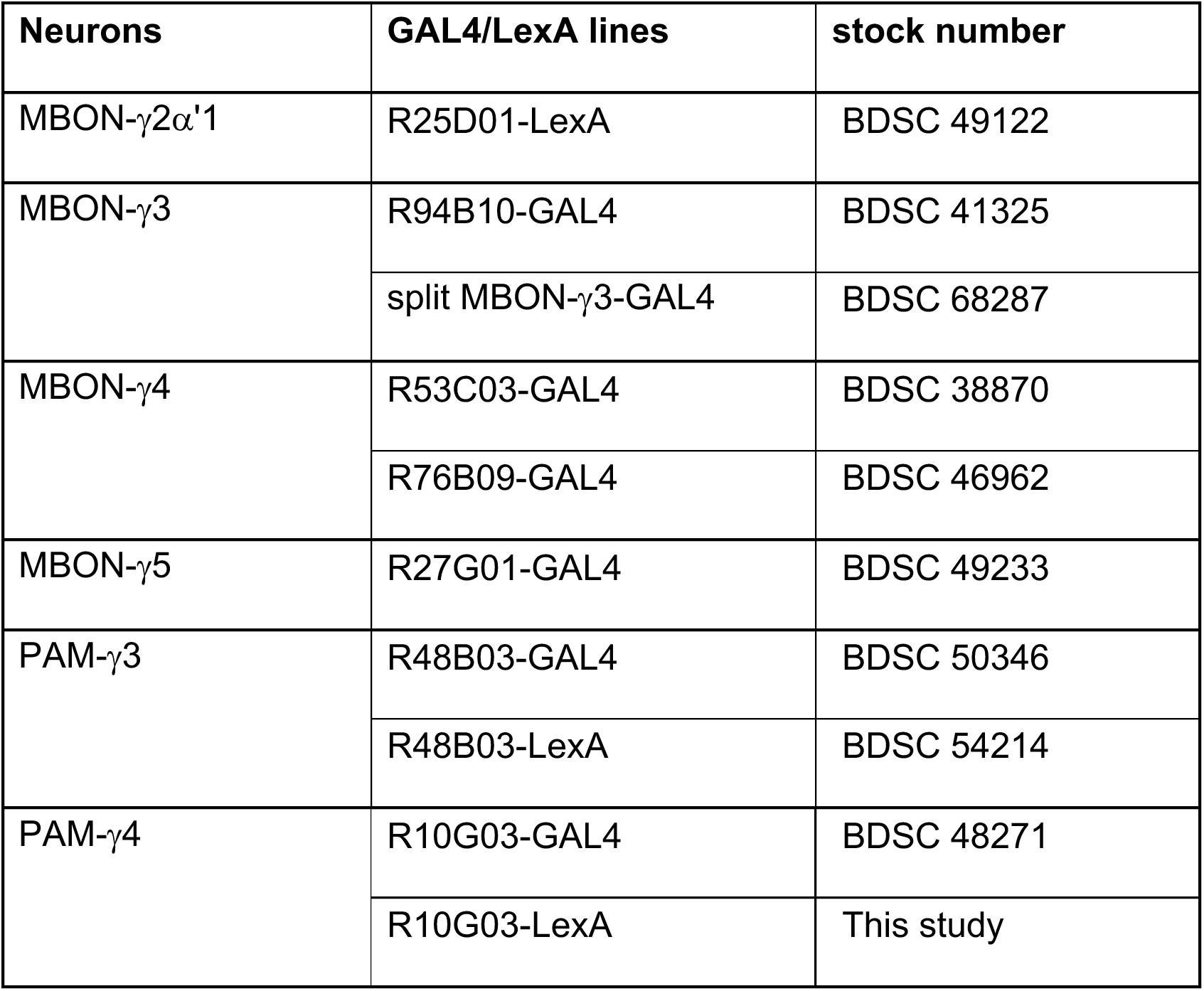
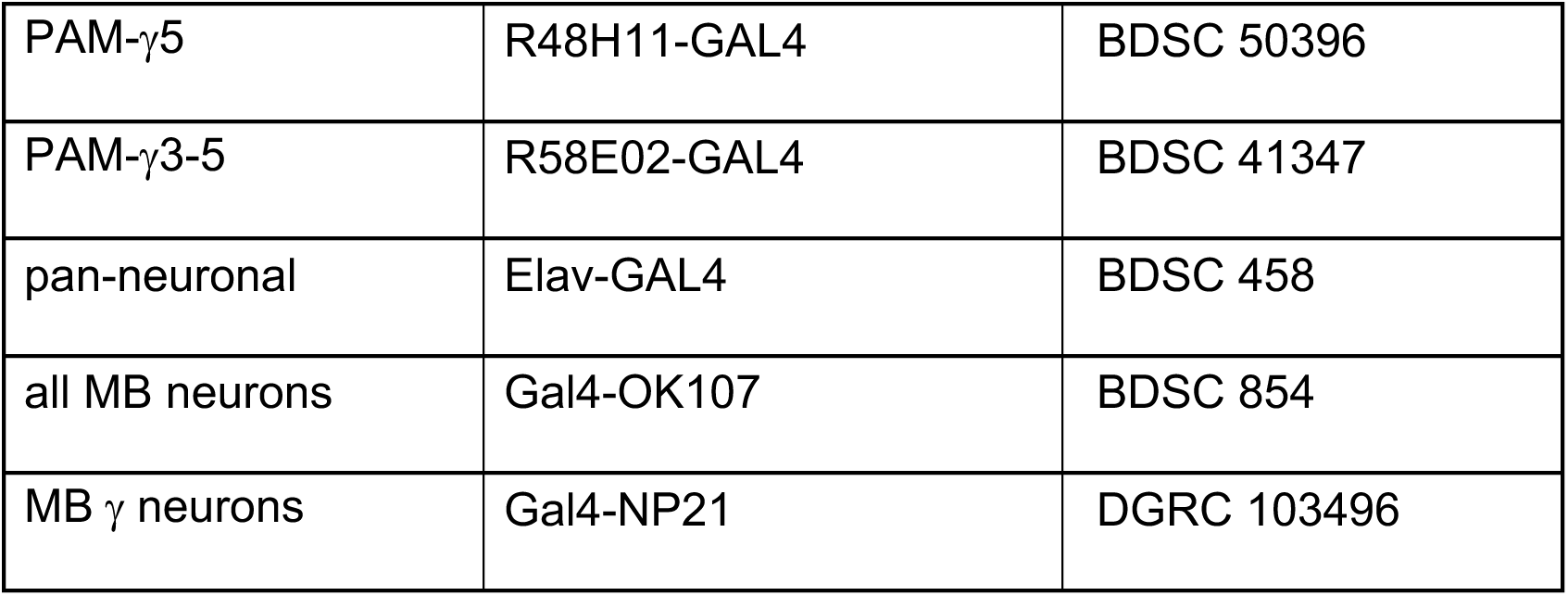
GAL4/LexA lines used for different types of neurons.

| Neurons | GAL4/LexA lines | stock number |
| --- | --- | --- |
| MBON-γ2α'1 | R25D01-LexA | BDSC 49122 |
| MBON-γ3 | R94B10-GAL4 | BDSC 41325 |
| | split MBON-γ3-GAL4 | BDSC 68287 |
| MBON-γ4 | R53C03-GAL4 | BDSC 38870 |
|  | R76B09-GAL4 | BDSC 46962 |
| MBON-γ5 | R27G01-GAL4 | BDSC 49233 |
| PAM-γ3 | R48B03-GAL4 | BDSC 50346 |
|  | R48B03-LexA | BDSC 54214 |
| PAM-γ4 | R10G03-GAL4 | BDSC 48271 |
|  | R10G03-LexA | This study |
| PAM-γ5 | R48H11-GAL4 | BDSC 50396 |
| PAM-γ3-5 | R58E02-GAL4 | BDSC 41347 |
| pan-neuronal | Elav-GAL4 | BDSC 458 |
| all MB neurons | Gal4-OK107 | BDSC 854 |
| MB $\gamma$ neurons | Gal4-NP21 | DGRC 103496 |

### Construction of R10G03-LexA plasmid and generation of transgenic lines

For constructing *R10G03-LexA plasmid*, the enhancer fragment R10G03 was amplified by PCR from genomic DNA with a forward primer 5’-GGGGACAAGTTTGTACAAAAAAGCAGGCTgccagcacactgcaactcctagagc, which contains the attB1 sequence in capital, and a reverse primer 5’-GGGGACCACTTTGTACAAGAAAGCTGGGTgatgaggcacttcagttgctgggaa, which contains the attB2 sequence in capital. The primers were designed based on the sequences available from the Janelia FlyLight website (https://flweb.janelia.org/cgi-bin/flew.cgi). The enhancer fragment containing the attB sequences was then cloned into the donor vector pDONR 211 (Invitrogen, Waltham, MA) at the attP1/attP2 sites using gateway BP Clonase II. The resulting entry clone was then used to transit the enhancer fragment to the pBPLexA::p65Uw destination vector using the LR clonase II. The constructs were injected into embryos of the *Drosophila* line y[1] w[*] P{y[+t7.7]=nanos-phiC31\int.NLS}X; PBac{y[+]-attP-9A}VK00027 (BDSC #35569) by the Rainbow Transgenic Flies, Inc. (Camarillo, California) and integrated into the VK27 attP docking site.

### Fluorescent immunostaining and Confocal microscopy imaging

*Drosophila* larval or adult brains were dissected and immunostained according to published protocol. Primary antibodies used include chicken anti-GFP (catalog #GFP-1020, Aves Labs, Tigard, Oregon; 1:500–1000), rabbit anti-DsRed (catalog #632496, Takara Bio USA, Inc., Mountain View, CA; 1:250), Chicken anti-spGFP1-10 (Abcam catalog # 13970, 1:1000), mouse anti-reconstituted spGFP1-10/spGFP11 (Sigma catalog# C6539, 1:100), mouse anti-Fas II monoclonal antibody (catalog #1D4, DSHB, 1:50). Secondary antibodies conjugated to Daylight 488 (1:100), Cy3 (1:500), Rhodamine Red-X (1:500), Daylight 647 (1:500) or Cy5 (1:500) used for immunostaining are from Jackson ImmunoResearch (West Grove, PA). Images were collected using a Carl Zeiss LSM780 confocal microscopy and processed with Adobe Photoshop.

### Quantifications and Statistical analyses

For quantifying the distance from the lateral or medial edges of the dendritic field of MBONs or axonal field of DANs to the midline of the brain, we measured the distance from the midpoint of the medial or lateral edges to the midline and we chose the focal slice in the middle of the z-stack for the measurement. For quantifying staining intensities of Eph-GFP in the γ3 or γ5 compartment in wild type or RNAi knockdown brains, the staining intensity was measured with the histogram in photoshop. The value was then subtracted by the background from areas with no GFP signals and normalized by the signal intensity (subtracted by the same background) in γ5 or γ3 compartment (i.e. quantification of the staining intensity in the γ3 compartment uses the staining intensity in the γ5 compartment for normalization, and *vice versa*). Student’s *t*-test was used for statistical analyses. Graphs were generated with Microsoft Excel.

## Supporting information

Ephrin_DengX_Supplemental Figures

## ACKNOWLEDGEMENTS

We thank Drs. R. Dearborn, J. Thomas, T. Chihara, S. Kunes, S. Rumpf for fly lines; the Bloomington *Drosophila* Stock Center and the TRiP at Harvard Medical School (NIH/NIGMS R01-GM084947), and Vienna *Drosophila* Resource Center for providing BAC and RNAi transgenic fly stocks; the Developmental Studies Hybridoma Bank for antibodies; members of the Zhu, Pignoni, and Lin Labs for thoughtful discussion and comments; Rainbow Transgenic Flies Inc for generating transgenic flies, Neuroscience Microscopy Core at Upstate Medical University for providing Zeiss LSM 780 confocal microscopy. This work was supported by the National Institute of Neurological Disorders and Stroke of the National Institutes of Health under Award Number R01NS085232 (S.Z.) and R21NS109748 (S.Z.).

## AUTHOR CONTRIBUTIONS

Conceptualization, X.D and S.Z.; Methodology; Investigation: X.D.; Data analysis: X.D., S.Z.; Writing-Original Draft, S.Z, Writing-Review & Editing-X.D. and S.Z.; Supervision, S.Z.; Funding, S.Z.

## DECLARE OF INTERESTS

The authors declare no competing interests.

