## Supplementary material for "Ephrin-mediated dendrite-dendrite repulsion regulates compartment-specific targeting of dendrites in the central nervous system": Ephrin_DengX_Supplemental Figures

Deng_Supplemental_Fig S1

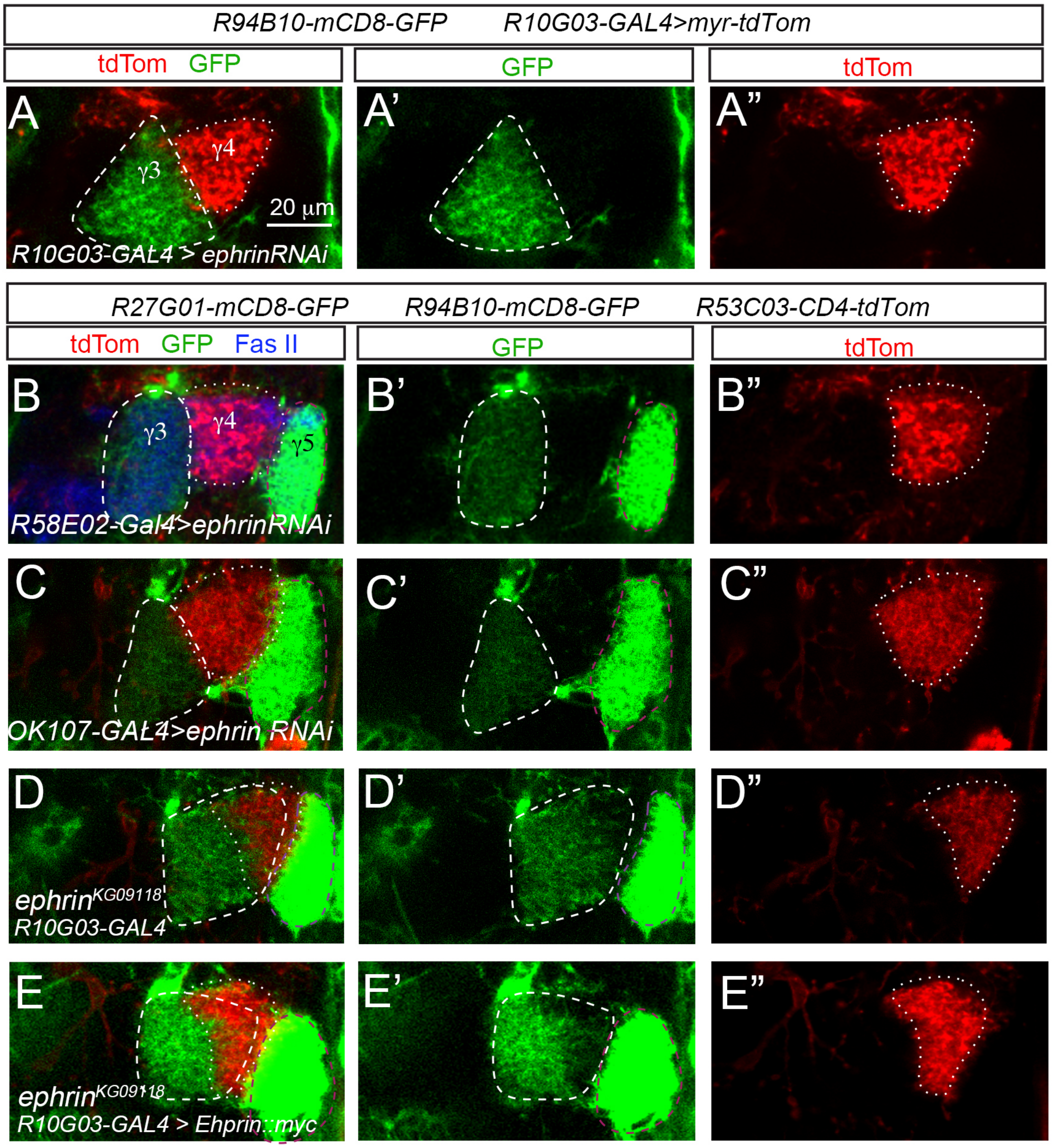

**Supplemental Fig. S1.** PAM-γ4 or MB neurons are not the source of Ephrin. In all images, MBON-γ3 dendrites are labeled with GFP and circled with white dashed lines, MBON-γ4 dendrites are labeled with tdTom and circled with white dotted lines, and MBON-γ5 dendrites are labeled with GFP and circled with purple dashed lines. (A-B') Projection of MBON-γ3 and MBON-γ4 neurons project their dendrites normally into the γ3 and γ4 compartment, respectively, when Ephrin is knocked down in PAM-γ4 neurons alone (A-A') or in PAM-γ3-5 neurons (B-B"). (C-C") MBON-γ3 dendrites project into the γ4 compartment and overlap with MBON-γ4 dendrites in *ephrin*$^{KG09118}$ homozygous mutant brains. (D-D") Expression of *UAS-Ephrin::myc* in PAM-γ4 neurons does not restores a sharp boundary between MBON-γ3 dendrites and MBON-γ4 dendrites in *ephrin*$^{KG09118}$ mutant brains. (E-E") knocking down Ephrin in MB neurons does not affect the compartment-specific targeting of the dendrites of MBON-γ3, MBON-γ4, and MBON-γ5 neurons.

Deng_Supplemental_Fig S2

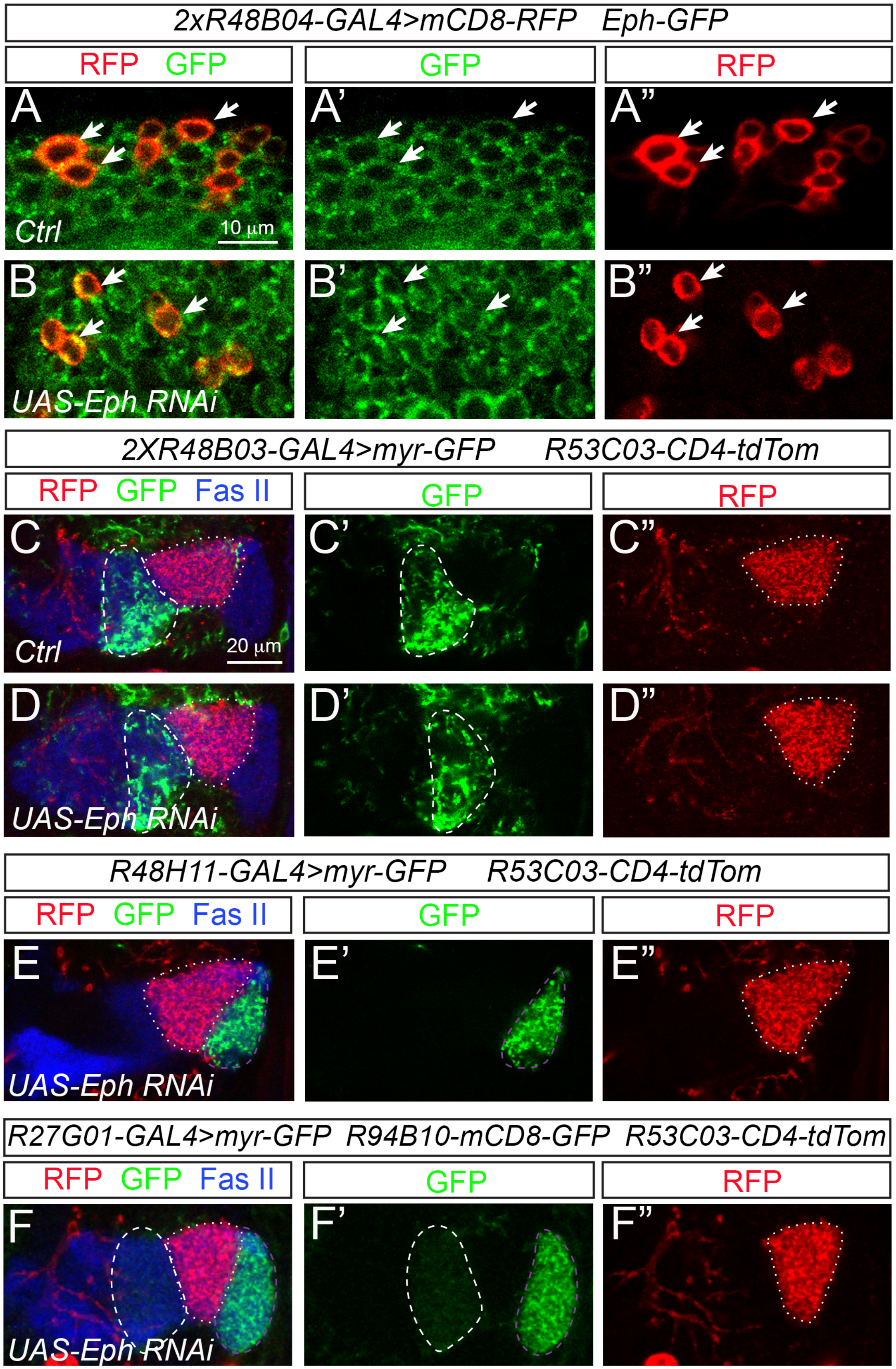

**Supplemental Fig. S2.** Eph is not required to restrict the projection of axons of PAM-γ3 and PAM-γ5 neurons and dendrites of MBON-γ5 neurons to their specific compartments. (A-A”) Eph-GFP expression in the cell bodies (arrows) of wild-type PAM-γ3 neurons labeled with RFP. (B-B”) Expression of *UAS-Eph RNAi* driven by even 2 copies of *R48B04-GAL4* in PAM-γ3 neurons does not reduce the Eph-GFP signal in their cell bodies (RFP, arrows). (C-C”) Projection of PAM-γ3 neuron axons (GFP, dashed circle) and MBON-γ4 dendrites (tdTom, dotted circles) in a wild-type adult brains. (D-D”) PAM-γ3 neuron axons (GFP, dashed circle) do not project into the γ4 compartment and overlap with MBON-γ4 dendrites (tdTom, dotted circles) when *UAS-Eph RNAi* is expressed in PAM-γ3 neurons. (E-E”) knockdown of Eph in PAM-γ5 neurons does not lead to projection of their axons (GFP, purple circles) into the γ4 compartment and overlap with MBON-γ4 dendrites (tdTom, dotted circles). (F-F”) knockdown of Eph in MBON-γ5 neurons does not affect the targeting of their dendrites (GFP, purple circles). They remain in the γ5 compartment and do not overlap with MBON-γ4 dendrites (tdTom, dotted circles). White dashed circles outline the dendritic area of MBON-γ5 neurons that are also labeled with GFP.

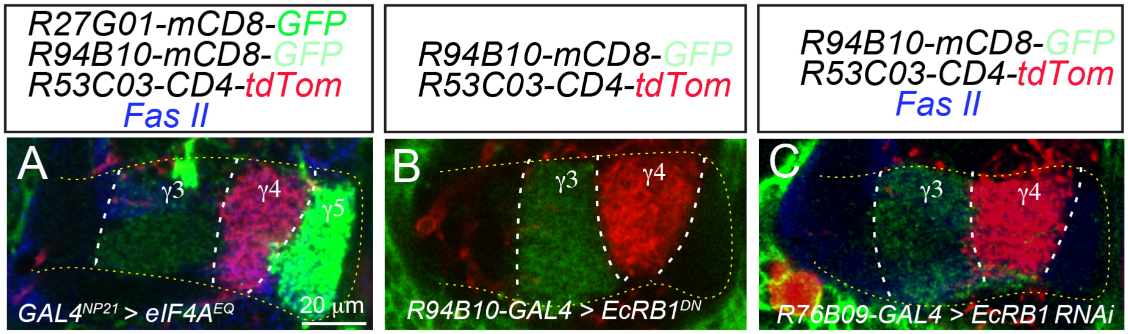


**Supplemental Fig. S3.** Remodeling is not involved in regulating compartment-specific targeting of MBON dendrites. In all images, MBON-γ4 neuron dendrites are labeled with tdTom, while MBON-γ3 dendrites and MBON-γ5 dendrites are labeled with weak and strong GFP expression, respectively. Fas II staining labels the MB γ lobe (outlined with yellow dotted lines). White dashed lines mark the boundaries between compartments. Dendrites of MBON-γ3, MBON-γ4, MBON-γ5 neurons are still projected into their respective compartments without overlapping with each other when $eIF4A^{EQ}$ (A), $EcRB1^{DN}$ (B), or EcRB1 RNAi (C) are expressed in the MB γ neurons, MBON-γ3 neurons, or MBON-γ4 neurons, respectively.

Deng_Supplemental_Fig S4

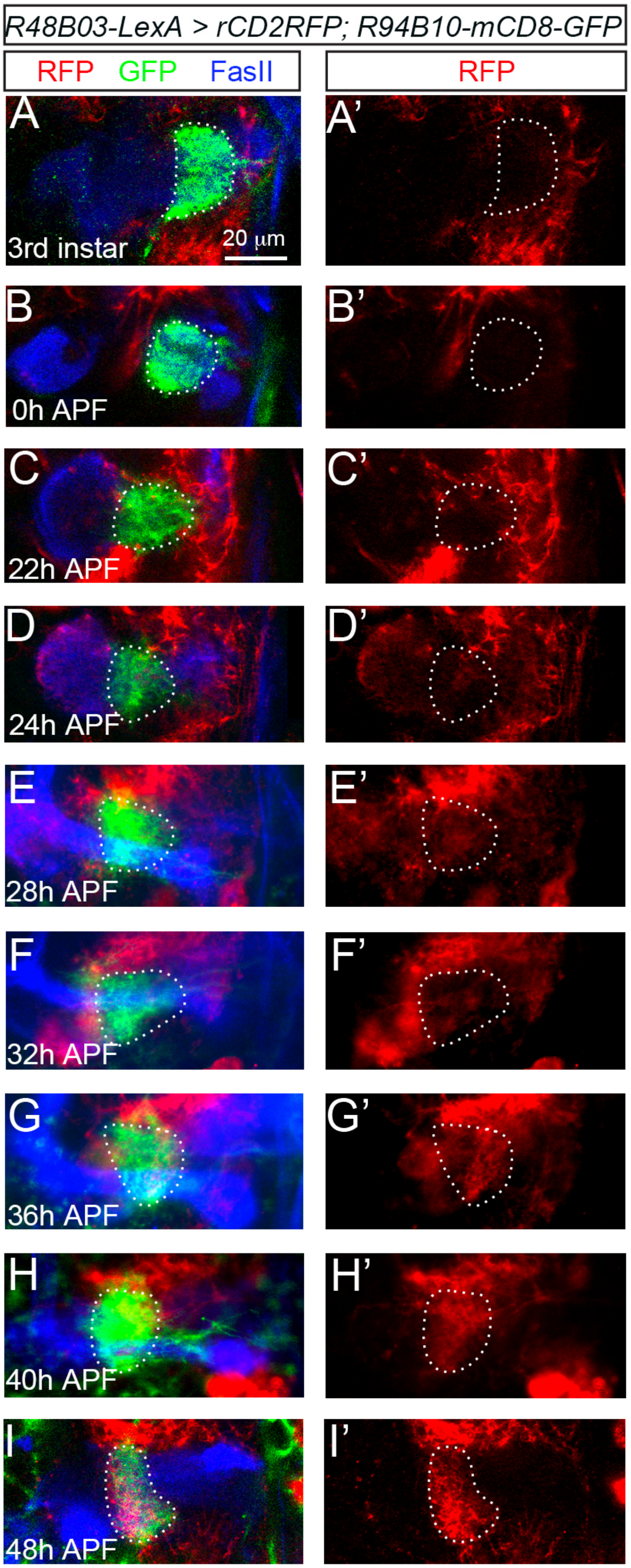

**Supplemental Fig. S4.** A time-course of the development of PAM-γ3 axons and MBON-γ3 dendrites. MBON-γ3 neurons are labeled with GFP and PAM-γ3 with RFP. MB axonal lobes are labeled with Fas II staining. Dotted circles outline the dendritic area of MBON-γ3 neurons. (A-B') No PAM-γ3 axons overlap with MBON-γ3 dendrites at the $3^{rd}$ instar larval stage (A-A') or 0 hrs APF (B-B'). (C-C') PAM-γ3 neurons start to project their axons into the area where MBON-γ3 dendrites reside at 22 hrs APF. (D-G') There is a gradual increase in the amount of PAM-γ3 axons that are projected into the area where MBON-γ3 dendrites reside from 24 to 40 hrs APF. (H-H') MBON-γ3 dendrites and PAM-γ3 axons adopt their projection patterns that are similar to those in adult brains at 48 hrs APF.
